# Investigating White Matter Functional Network Connectivity Across the Alzheimer’s Disease Spectrum Using Resting-State fMRI

**DOI:** 10.64898/2026.02.04.703913

**Authors:** Vaibhavi S. Itkyal, Theodore J. LaGrow, Kyle M. Jensen, Armin Iraji, Vince D. Calhoun

## Abstract

White matter (WM) has traditionally been considered structurally important but functionally inert in fMRI research. However, growing evidence indicates that WM exhibits meaningful BOLD fluctuations and participates in functional connectivity. Here, we investigate alterations in WM functional network connectivity (FNC) across the Alzheimer’s disease (AD) spectrum using resting-state fMRI data from the Alzheimer’s Disease Neuroimaging Initiative (ADNI; 415 cognitively normal (CN), 283 mild cognitive impairment (MCI), 91 AD). We applied a guided independent component analysis (ICA) approach based on a combined multiscale template including 202 intrinsic connectivity networks (ICNs; 97 WM, 105 gray matter (GM)) to estimate subject-specific timecourses and compute static FNC (sFNC). Group differences in WM–WM, GM–GM, and WM–GM connectivity (AD–CN, AD–MCI, MCI–CN) were assessed using two-sample t-tests with covariates for age, sex, and motion, with false discovery rate correction. Results showed robust alterations in WM–WM and WM–GM connectivity in AD, particularly involving WM subcortical, frontal, sensorimotor, and occipitotemporal networks. Several WM–GM interactions with cerebellar and hippocampal GM networks were also disrupted, including reduced GM–cerebellar:WM–frontal coupling and increased GM–hippocampal:WM– frontal connectivity. Notably, MCI already showed WM–GM dysconnectivity relative to CN, suggesting that functional disruption of WM circuits emerges prior to overt dementia. These findings provide converging evidence that WM functional connectivity is both measurable and selectively altered across the AD continuum. Our findings support WM sFNC as a complementary candidate biomarker to GM-based measures for staging and monitoring AD. This is, to our knowledge, the first large-scale ADNI study to jointly model WM and GM intrinsic connectivity networks and quantify WM–GM dysconnectivity across CN, MCI, and AD.

## Introduction

Alzheimer’s disease (AD) is a progressive neurodegenerative disorder characterized by memory impairment, cognitive decline, and large-scale disruption of brain networks. While most functional MRI (fMRI) studies have focused on gray matter (GM) pathology and connectivity, converging evidence suggests that white matter (WM) also plays a critical role in AD progression (Bubb, Metzler-Baddeley, and Aggleton 2018; Kaskikallio et al. 2019). Diffusion imaging studies consistently report microstructural abnormalities in WM, including myelin degradation and axonal loss, across the AD continuum (Mito et al. 2018). Yet, the functional contribution of WM to AD-related network changes remains comparatively underexplored.

Historically, WM has been largely neglected in fMRI analyses because of its lower blood-oxygenation level-dependent (BOLD) signal amplitude and concerns about vascular and partial-volume effects. Recent work, however, has shown that WM exhibits structured, low-frequency BOLD fluctuations that are temporally coherent, spatially organized, and reproducible across datasets and sites (Peer et al. 2017; Gore et al. 2019). These findings challenge the view of WM as functionally “silent” and instead support the idea that WM participates actively in large-scale communication between GM regions.

Independent component analysis (ICA) has been particularly useful for characterizing WM functional organization. ICA-derived intrinsic connectivity networks (ICNs) have been identified within WM that are distinct from GM networks and show stable patterns across individuals and cohorts (Itkyal et al. 2025). Such WM ICNs can be grouped into functional subdomains (e.g., sensorimotor, occipitotemporal, frontal, subcortical), providing a systems-level framework to study WM functional involvement in health and disease. In parallel, extensive work on GM ICNs has revealed consistent alterations in default mode, hippocampal, frontal, and sensorimotor networks in AD and mild cognitive impairment (MCI), the intermediate clinical stage of the disease (LaGrow et al. 2025).

Functional network connectivity (FNC) computed from ICN time course offers a compact summary of large-scale interactions among brain networks. Static FNC (sFNC), obtained from pairwise correlations over the full scan, has been widely used to identify modular alterations in GM connectivity in AD. However, most existing studies focus exclusively on GM or treat WM as nuisance signal, leaving open the question of how WM functional connectivity and WM–GM interactions change across the AD continuum. A joint analysis of WM and GM sFNC could provide a more complete picture of network degradation and reorganization in AD, and may reveal patterns that are not apparent from GM-only analyses.

In this brief report, we leverage a guided ICA framework that combines a large-scale, validated WM ICN template (97 networks freely available at https://trendscenter.org/data/) with a GM network template (105 networks) (Iraji et al. 2023, Jensen et al. 2024) to estimate subject-specific ICNs and whole-brain sFNC in a large Alzheimer’s Disease Neuroimaging Initiative (ADNI) cohort. Using resting-state fMRI data from cognitively normal (CN), MCI, and AD participants, we examine group differences in WM–WM, GM–GM, and WM–GM connectivity across the disease spectrum. We hypothesized that: (i) WM–WM and WM–GM sFNC would show robust alterations in AD relative to CN, particularly in posterior and subcortical-frontal pathways implicated in AD; and (ii) WM-related dysconnectivity would be observed even at the MCI stage of disease progression, indicating that WM functional changes emerge prior to overt dementia. By explicitly incorporating WM into a network-level framework, this work aims to evaluate the potential of WM functional connectivity as a complementary imaging biomarker for Alzheimer’s disease.

## Methods

### Participants

This study utilized resting-state functional magnetic resonance imaging (rs-fMRI) data from the Alzheimer’s Disease Neuroimaging Initiative (ADNI) (Mueller et al. 2005). The sample included 789 participants spanning the Alzheimer’s disease spectrum: 415 cognitively normal (CN), 283 individuals with mild cognitive impairment (MCI), and 91 individuals with a clinical diagnosis of Alzheimer’s Disease (AD). Diagnostic classification was based on ADNI consensus criteria, incorporating clinical evaluation, cognitive testing, and informant reports. Participants were included if they had a usable rs-fMRI scan, corresponding high-resolution T1-weighted anatomical image, and complete demographic information. All data were acquired on 3T MRI systems at ADNI sites using standardized protocols; detailed acquisition parameters are available in ADNI documentation and are summarized in the Supplement. Demographic characteristics for each diagnostic group (including age and sex distribution) are listed in Table 1. Subject inclusion criteria and clinical diagnoses followed ADNI protocols, and all procedures were approved by local institutional review boards with informed consent obtained from each participant.

**Table 1.** Group-wise summary statistics for cognitively normal (CN), mild cognitive impairment (MCI), and Alzheimer’s disease (AD) participants.

| Research Group | # Scans (F/M) | Age, mean ± SD (years) |
| --- | --- | --- |
| CN | 415 (247/168) | 71.20±6.45 |
| MCI | 283 (126/157) | 71.82±7.38 |
| Dementia | 91 (45/46) | 73.48±7.80 |

### fMRI Preprocessing

All rs-fMRI data underwent standard preprocessing using a combination of SPM12 and FSL software (following the procedures outlined in the NeuroMark framework (Du et al. 2020)). The first five volumes of each scan were discarded to allow for magnetization stabilization. The remaining volumes were then realigned using rigid-body motion correction to estimate and correct for head movement across the time series. Slice timing correction was applied to adjust for inter-slice acquisition delays relative to the middle slice.

Following motion and slice-timing correction, functional images were spatially normalized to the Montreal Neurological Institute (MNI) standard space using deformation fields estimated from each subject’s T1-weighted anatomical image. Normalized images were resampled to 3 mm^3^ isotropic voxels to provide a common resolution across participants. A Gaussian spatial smoothing kernel of 6 mm full-width at half maximum (FWHM) was then applied to improve signal-to-noise ratio while maintaining a balance between spatial specificity and sensitivity in both WM and GM.

To ensure data quality and minimize motion-related confounds, we implemented stringent exclusion criteria at the subject level. Scans were discarded if they contained fewer than 120 usable time points, if the mean framewise displacement exceeded 0.25 mm, or if absolute head motion surpassed 3 mm of translation or 3 degrees of rotation in any direction. After applying these quality control thresholds, more than 700 scans across the three diagnostic groups (CN, MCI, AD) were retained for subsequent functional network and connectivity analyses.

### WM and GM Functional Network Templates and Guided ICA

To characterize brain-wide functional organization, we utilized a previously validated WM ICNs template comprising 97 WM components. These WM ICNs were derived from a large cohort of healthy individuals using spatial ICA (A. Iraji et al. 2023), and were selected based on their spatial specificity, reproducibility, and low overlap with GM and CSF. The WM ICNs were grouped into functional subdomains (e.g., sensorimotor, occipitotemporal, frontal, subcortical, parietal), providing a systems-level representation of WM functional architecture. Details of the template derivation and subdomain labels are provided in prior work (Itkyal et al. 2025) and summarized in the Supplement.

To model GM functional organization in parallel, we incorporated 105 ICNs from an established GM functional network template (A. Iraji et al. 2023, Jensen et al. 2024). These GM components include canonical resting-state networks such as default mode, hippocampal/medial temporal, sensorimotor, visual, cerebellar, and fronto-parietal systems. Together, the WM (97) and GM (105) templates yielded a combined set of 202 spatial priors that span both tissue types.

Subject-specific ICNs and their associated timecourses were estimated from each subject using a multi-objective optimization ICA with reference (MOO-ICAR) framework. This guided ICA approach imposes soft spatial constraints based on the template ICNs while capturing individual variability. In brief, for each subject, the algorithm optimizes a cost function that simultaneously maximizes statistical independence of components and spatial similarity to the WM and GM template maps. This yields subject-level ICN spatial maps that closely align with the group priors, along with corresponding timecourses that reflect the subject’s spontaneous BOLD fluctuations in each network.

By applying MOO-ICAR with the combined WM and GM template, we obtained 202 subject-specific ICNs per participant (97 WM, 105 GM), each with a single representative timecourse. These timecourses formed the basis for subsequent functional connectivity analyses of WM–WM, GM–GM, and WM–GM interactions.

Spatial maps of the (A) 97 WM ICNs and (B) 105 GM ICNs used as priors in the guided ICA framework (see **Figure 1**). WM ICNs are grouped into 13 functional subdomains (e.g., sensorimotor, occipitotemporal, frontal, subcortical, parietal), and GM ICNs are grouped into 14 canonical resting-state subdomains (e.g., default mode, hippocampal/medial temporal, sensorimotor, visual, cerebellar, fronto-parietal). ICN maps are displayed in MNI space and thresholded for visualization. These templates provide the spatial priors for estimating subject-specific ICNs spanning both WM and GM.

**Figure 1.**
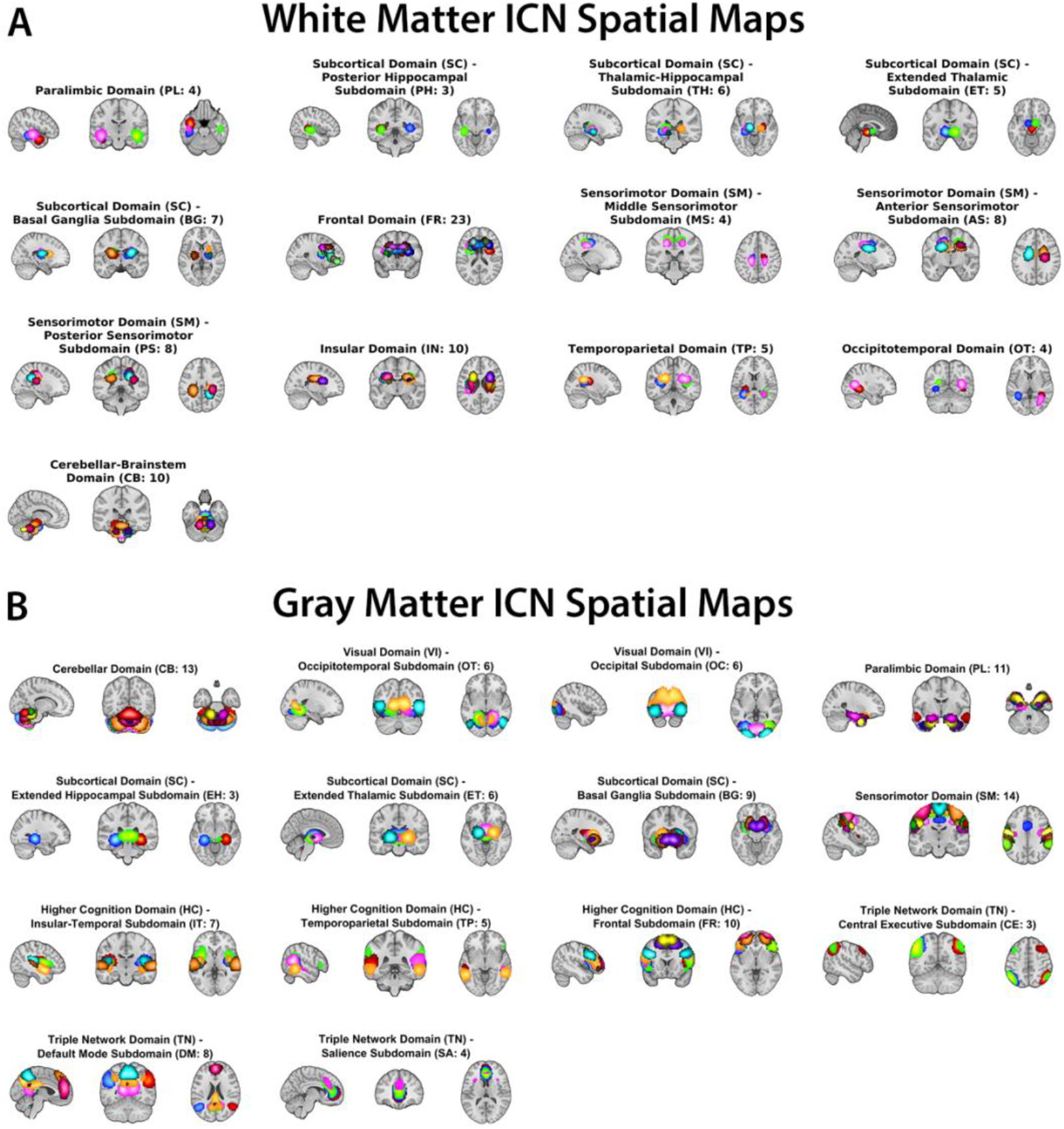
White matter (A, top) and gray matter (B, bottom) intrinsic connectivity network (ICN) templates.

### Functional Connectivity and Statistical Analysis

Static FNC was computed from the ICN timecourses for each subject. Specifically, we calculated pairwise Pearson correlations between all 202 ICN timecourses (97 WM + 105 GM), yielding a 202 × 202 symmetric connectivity matrix per participant. Correlation coefficients were transformed to Fisher’s z-scores to improve normality, producing whole-brain sFNC profiles for each individual.

For descriptive purposes, sFNC edges were categorized into three classes based on the tissue type of the participating ICNs: (i) WM–WM: connections between two WM ICNs,(ii) GM–GM: connections between two GM ICNs, and (iii) WM–GM: cross-tissue connections between a WM ICN and a GM ICN.

Group-level mean sFNC matrices were computed separately for CN, MCI, and AD to visualize overall network organization and qualitative differences across the disease spectrum. To formally assess group differences, we performed two-sample t-tests on each sFNC edge for three contrasts: AD vs. CN, AD vs. MCI, and MCI vs. CN. All models included age, sex, and mean framewise displacement as covariates to account for demographic and motion-related effects.

Multiple comparisons were controlled using false discovery rate (FDR) correction at q < 0.05 across all tested edges within each contrast. For visualization, significant edges were displayed in difference matrices using a signed –log10(p) scaling multiplied by the sign of the t-statistic, highlighting both the strength and direction of connectivity changes. To summarize the distribution of effects, we additionally tabulated the number and proportion of significant WM–WM, GM–GM, and WM–GM connections for each contrast. More detailed, subdomain-level summaries of sFNC changes are reported in the Results and Supplementary Material.

## Results

### Preserved large-scale WM and GM network organization across groups

sFNC matrices showed a broadly similar large-scale organization across CN, MCI, and AD groups (Figure 2). In all three diagnostic categories, the combined WM and GM network representation exhibited a clear block structure corresponding to WM–WM, GM– GM, and WM–GM connections, as well as modular organization within sensorimotor, visual, cerebellar, frontal, and subcortical systems. This preserved global architecture indicates that core network modules remain identifiable across the AD spectrum, despite substantial reconfiguration of specific within- and cross-tissue connections described below.

**Figure 2:**
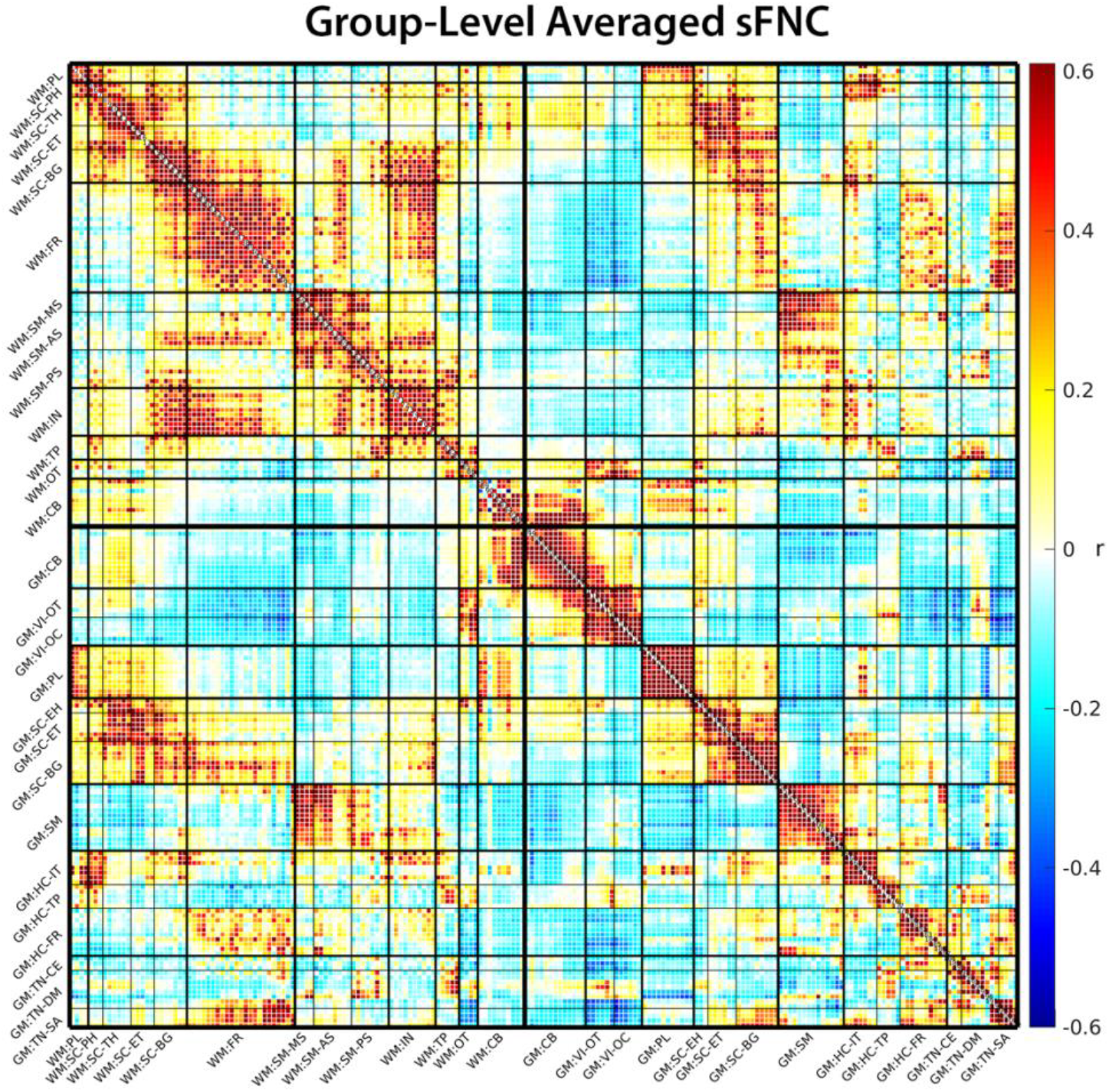
Group-level averaged sFNC matrix. The sFNC matrix is modularized into 13 WM subdomains and 14 GM subdomains, indicated with a ‘WM:’ or ‘GM:’ prefix respectively. The group-average sFNC shows clear modular organization in both WM and GM subdomains. WM subdomains form comparatively broader modules, while GM exhibits finer-grained modular structure. Notable WM–GM coupling is evident, particularly involving subcortical, frontal, and somatosensory subdomains.

### Widespread white matter and cross-tissue connectivity disruption in Alzheimer’s disease

Group comparisons across the AD spectrum revealed widespread alterations in functional connectivity involving both WM and GM networks. These disruptions were particularly evident in contrasts involving the AD group, AD versus CN and AD versus MCI, highlighting the involvement of WM and WM–GM interactions in disease progression.

In the AD versus CN comparison (Figure 3A), we observed distinct alterations in WM– WM connectivity. Hypoconnectivity emerged within the occipitotemporal (OT) WM subdomain and in its connections to other WM networks, suggesting reduced integration across posterior WM pathways. In contrast, hyperconnectivity was evident between subcortical (SC) and sensorimotor (SM) WM domains, potentially reflecting compensatory or inefficient reorganization of subcortical–sensorimotor communication in AD.

**Figure 3.**
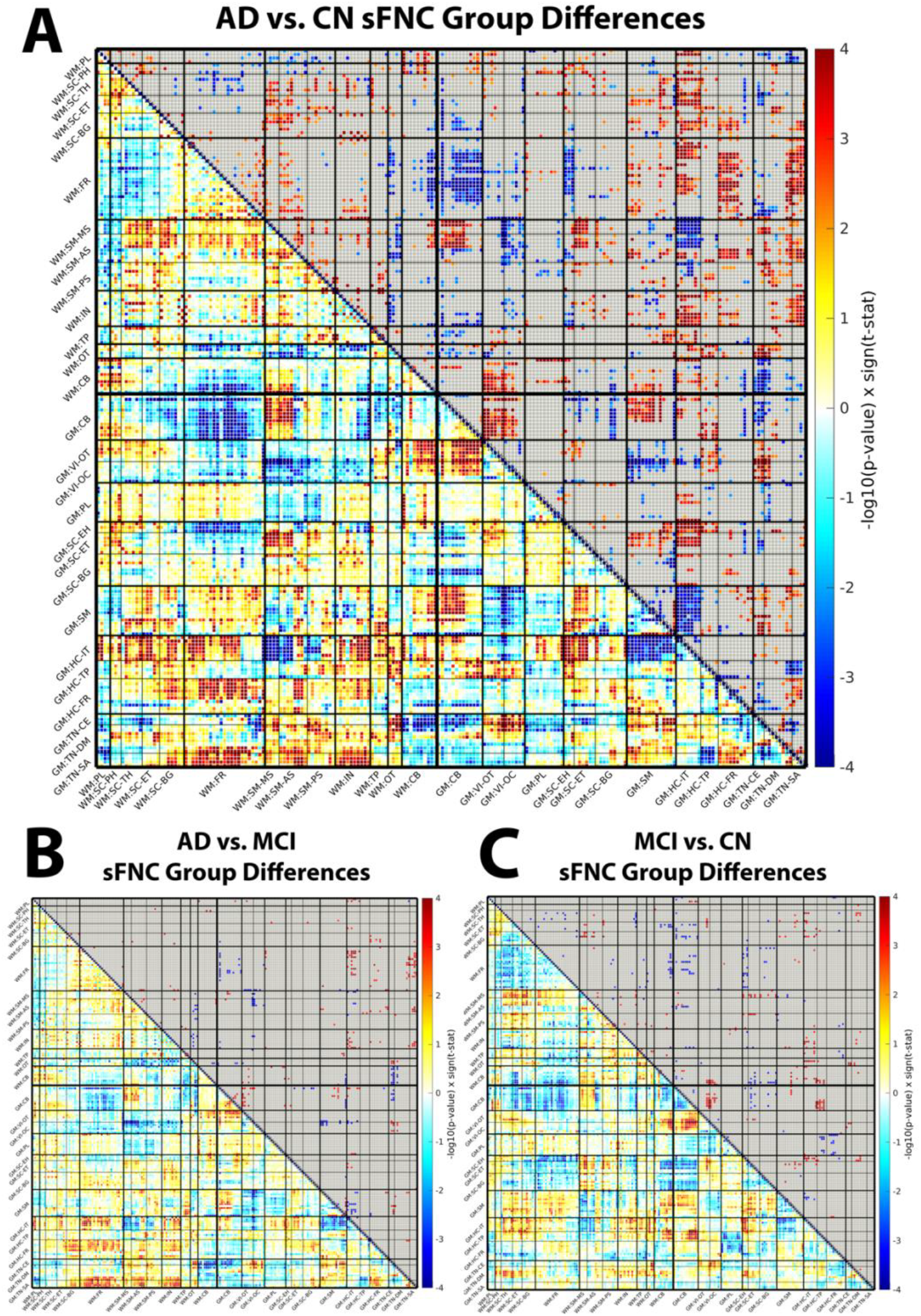
Group differences in static functional network connectivity (sFNC) across diagnostic stages. Difference matrices for (A) AD vs. CN, (B) AD vs. MCI, and (C) MCI vs. CN contrasts. Each matrix summarizes edge-wise group differences in sFNC between 202 ICNs (97 WM, 105 GM), ordered by WM and GM subdomains as in Figure 2. Color values represent signed –log_10_(p) multiplied by the sign of the corresponding t-statistic from two-sample t-tests, with age, sex, and mean framewise displacement as covariates. Positive (red) values indicate higher connectivity in the first group (e.g., AD) relative to the second (e.g., CN), while negative (blue) values indicate lower connectivity. Edges are thresholded at a false discovery rate (FDR)–corrected q < 0.05. Together, these matrices highlight widespread disruptions in WM–WM, GM–GM, and WM–GM connectivity, including prominent posterior WM hypoconnectivity, subcortical–sensorimotor reorganization, and early cerebellar/subcortical–frontal WM–GM dysconnectivity across the AD spectrum.

Within the GM–GM domain, significant hypoconnectivity was observed between SM and hippocampal (HC) regions, implicating disrupted interactions between motor-related and memory-related systems. At the same time, enhanced connectivity between cerebellar (CB) and visual (VI) subdomains suggested atypical reorganization within visuomotor circuits.

Critically, several WM–GM interactions were also disrupted in AD relative to CN. These included reduced connectivity between GM–CB and WM–frontal (FR) networks, alongside increased coupling between GM–HC and WM–FR regions. Together, these patterns highlight altered communication between memory-related GM regions, cerebellar systems, and frontal WM pathways, indicating that AD-related network disruption is not confined to GM but extends prominently to cross-tissue interactions.

### Progressive reorganization of white matter–gray matter connectivity from MCI to AD

In the AD versus MCI comparison (Figure 3B), differential connectivity patterns continued to emerge, indicating progressive network reorganization beyond the MCI stage. At the WM–GM level, we observed increased connectivity between GM temporal and semantic subdomains, such as temporal pole and superior temporal sulcus regions, and WM–FR and WM–insular (IN) networks. These findings suggest that temporal and semantic GM systems become more tightly coupled to frontal and insular WM pathways as disease severity increases.

Concurrently, several GM–GM and WM–GM connections showed reduced connectivity in AD relative to MCI. Decreases were evident between GM–SM and GM–HC/insular– temporal (HC–IT) regions, as well as between GM–VI and WM–SM networks. WM–WM integration was also reduced, particularly between OT and SM domains, pointing to ongoing disconnection of posterior and sensorimotor WM pathways even between adjacent clinical stages. This combination of strengthened temporal–frontal/insular cross-tissue coupling and weakened sensorimotor and posterior pathways is consistent with a progressive, spatially nonuniform reorganization of both within-tissue and cross-tissue networks from MCI to AD.

### Early white matter–gray matter dysconnectivity in mild cognitive impairment

The MCI versus CN contrast (Figure 3C) revealed that WM–GM dysconnectivity is already present at the prodromal stage. Widespread reductions in WM–GM connectivity were most prominent between GM–CB and GM–SC regions and WM–FR subdomains, suggesting that cerebellar and subcortical influences on frontal WM pathways are diminished before the onset of overt dementia. Within the GM–GM domain, we observed reduced CB–CB connectivity accompanied by increased CB–VI coupling. This opposing pattern, loss of intra-cerebellar connectivity with heightened cerebellar–visual interactions, may reflect early compensatory processes aimed at maintaining functional stability in the face of emerging pathology.

Collectively, these results emphasize the evolving nature of WM involvement in AD. Altered connectivity within and across WM and GM domains from CN to MCI to AD supports the view that WM plays an active, dynamic role in the neurodegenerative process. These functional disruptions extend beyond structural degeneration alone and highlight the potential of WM-based functional connectivity measures as complementary biomarkers for early diagnosis and disease monitoring. Across contrasts, cerebellar/subcortical–frontal WM–GM connectivity appears to be altered earliest (MCI vs. CN), with subsequent disruption of posterior OT and sensorimotor WM–WM connectivity and broader WM–GM reweighting in AD.

## Discussion

This study provides evidence that WM functional connectivity is altered across the AD spectrum and that these alterations are not restricted to GM networks. By jointly modeling WM and GM ICNs, we identified stage-dependent disruptions in sFNC within WM, within GM, and across WM–GM connections in CN, MCI, and AD groups.

A consistent pattern across contrasts was reduced connectivity in posterior WM pathways, particularly within occipitotemporal (OT) subdomains, together with increased connectivity between subcortical (SC) and sensorimotor (SM) WM networks. The posterior hypoconnectivity is compatible with prior diffusion MRI reports of microstructural degradation in posterior association and visual tracts in AD (Mito et al., 2018) and suggests weakened functional coordination in these pathways. In contrast, SC–SM hyperconnectivity may index secondary reorganization of subcortical–sensorimotor loops, consistent with prior observations of altered subcortical–cortical coupling in AD (J. Zhao et al. 2019).

At the GM level, we observed decreased connectivity between sensorimotor (SM) and hippocampal (HC) subdomains, indicating decoupling between memory-related and sensorimotor systems, in line with known hippocampal and default-mode disruptions in AD (F. Zhao et al. 2008). In addition, altered cerebellar (CB) connectivity, including reduced CB–CB coupling and increased CB–visual (VI) interactions, suggests a shift from intra-cerebellar coordination toward cerebellar–cortical and cerebellar–visual engagement. Prior work has implicated cerebellar contributions to visuomotor, navigational, and cognitive functions in aging and dementia (Jacobs et al. 2018), and the observed pattern is consistent with a redistribution of functional load across cerebellar and cortical systems.

Cross-tissue analyses indicated that WM–GM connections are affected early and evolve with disease severity. In AD, reduced connectivity between GM–CB/SC regions and WM– frontal (FR) networks, together with increased connectivity between GM–HC and WM– FR, indicates abnormal reweighting of inputs to frontal executive pathways from cerebellar, subcortical, and hippocampal systems. These findings generalize earlier GM-only reports of disrupted fronto-hippocampal coupling in AD (Wang et al. 2006; Li et al. 2021) by showing that part of this disruption is expressed in WM–GM interfaces. Importantly, WM–GM dysconnectivity was already present in MCI relative to CN, especially in cerebellar/subcortical–frontal connections, and continued to change between MCI and AD. This pattern is consistent with the broader literature describing coexisting hypo- and hyperconnectivity and network instability in AD (Jones et al. 2016; Dai et al. 2019).

These results support the view that WM functional connectivity provides information beyond structural metrics alone. Structural imaging robustly demonstrates WM degradation in AD (Mito et al. 2018), but does not quantify how communication across distributed systems is functionally altered. By estimating sFNC from WM and GM ICNs, we capture how the system-level organization of WM and WM–GM interactions changes with disease. This aligns with work showing that WM exhibits reliable, low-frequency BOLD fluctuations that are temporally coherent and spatially organized (Peer et al. 2017; Gore et al. 2019; Itkyal et al. 2025), and extends it by demonstrating that these fluctuations carry disease-relevant, stage-specific information.

Several limitations should be noted. First, the present analysis is restricted to static FNC; dynamic FNC methods may reveal time-varying connectivity states that are more sensitive to early or subtle abnormalities. Second, although the use of validated WM and GM ICN templates and a guided ICA framework improves reproducibility, template-based approaches can still be sensitive to residual noise, individual anatomical variability, and potential template–subject mismatch. Third, the design is cross-sectional and does not include behavioral, cognitive, or molecular (e.g., amyloid, tau) measures, limiting inference about temporal trajectories and clinical correlates of the observed connectivity changes. Finally, multi-site acquisition in ADNI introduces scanner and site heterogeneity that may not be fully accounted for by standard preprocessing and covariate adjustment.

Future work should examine WM and WM–GM connectivity using dynamic FNC approaches, integrate diffusion MRI to relate structural and functional alterations within specific tracts, and incorporate cognitive and biomarker measures in longitudinal designs to test whether WM-related connectivity metrics predict conversion or rate of decline. Task-based fMRI could further clarify how WM–GM connectivity differences modulate performance in memory, visuospatial, or motor tasks. In addition, a direct comparison of structural connectivity in WM with these WM FNC results may yield additional information about how brain structure and brain function are jointly and differentially impacted by neurodegenerative disease (Majeed et al., 2011, LaGrow et al., 2026, Bolt et al., 2022, Liu et al., 2018).

In summary, this study shows that WM functional network connectivity can be robustly quantified in standard rs-fMRI data and that it is systematically altered from CN to MCI to AD. Joint analysis of WM and GM networks reveals widespread disruptions in intra- and cross-tissue connectivity and identifies WM–GM connections that change early and progress with disease severity. These findings support the incorporation of WM-based functional measures into connectivity frameworks for Alzheimer’s disease and motivate evaluation of WM-inclusive sFNC features as complementary biomarkers for diagnosis, stratification, and monitoring.

## Ethics statement

This study used de-identified data from the Alzheimer’s Disease Neuroimaging Initiative (ADNI). All ADNI participants provided written informed consent, and study protocols were approved by the institutional review boards of each participating site.

## Data availability statement

The datasets analyzed in this study are publicly available from the Alzheimer’s Disease Neuroimaging Initiative (ADNI) at adni.loni.usc.edu. Access to raw imaging and clinical data requires registration and adherence to ADNI data use agreements. Derived functional connectivity matrices and analysis code are available from the corresponding author on reasonable request.

## Author contributions

VSI conceived and designed the study, performed data preprocessing, conducted all analyses, carried out post-processing, generated visualizations and wrote the first draft of the manuscript. TJL and VDC contributed to the interpretation of results and helped revise and refine all subsequent versions of the manuscript. AI and KMJ offered methodological guidance and critical revisions. The project was completed under the supervision of VDC. All authors approved the final manuscript.

